# Cross-chemical and cross-species toxicity prediction: benchmarking and a novel 3D-structure-based deep learning model

**DOI:** 10.1101/2025.11.24.690199

**Authors:** Ruying Yuan, Joseph Shaw, Haixu Tang, Yuzhen Ye

## Abstract

Prediction of a compound’s toxicity is a key step toward realizing animal-free testing of chemical compounds. Recent advances have yielded significant progress in computational toxicity prediction, including machine learning methods that utilize chemical fingerprints and deep learning-based latent representations. However, challenges remain, primarily due to the lack of clean training datasets and the inconsistent model performance. To address these challenges, we curated a comprehensive dataset of aquatic toxicity from seven data sources, which contains 50,603 records for 5,889 compounds across 2,285 different species, much larger than similar datasets used in previous studies. We also developed *tox-learn*, a Python library featuring tools for automated dataset cleaning, machine learning methods and performance evaluation. The library places special emphasis on avoiding overestimation of prediction accuracy caused by improper train-test data splitting. Based on this toolbox, we benchmarked various predictive models using different train-test splitting strategies on the curated dataset. Our results showed that the choice of machine learning method, molecular fingerprint, and train-test splitting strategy all significantly affect performance. We demonstrated that incorporating species information generally improved predictions, although the degree of improvement depended on how this information was represented. In addition, we developed a new 3D structure–based deep-learning model, *3DMol-Tox*, which achieves regression accuracy comparable to the best 2D-structure based model (GPBoost) while exhibiting consistently higher within-one-bin (W1B) classification accuracy. Finally, we analyzed the impact of different train–test splitting strategies and provide recommendations based on our benchmarking, such as using structure-aware splitting to mitigate information leakage, a common issue that inflates reported model performance.

## Introduction

Predicting the toxicity of chemical compounds to animals and humans is critical for protecting environmental and human health, and for advancing efforts to reduce reliance on animal testing. Over the past decade, *in silico* approaches have emerged as promising alternatives to traditional animal-based toxicity tests. Methods such as quantitative structure–activity relationships (QSARs), read-across structure–activity relationships (RASARs), and deep learning–based generative models have demonstrated the potential to predict toxicity from chemical structure and related data. For example, Bio-QSARs [30] integrate chemical descriptors and physiological traits, enabling cross-species and cross-chemical predictions. Approaches such as RASARs [19] use binary fingerprints and chemical similarity to predict hazard profiles, while more recent generative models like Tox-GAN [8] and AnimalGAN [7] can simulate complex biological responses, including gene expression and clinical pathology outcomes.

Despite these advances, major challenges persist. A critical limitation is the lack of high-quality, harmonized datasets that accurately represent chemical diversity and species variability while minimizing data leakage and bias. In some studies, such as the original Bio-QSAR work, the presence of identical chemical structures in both training and test sets has led to artificially inflated performance metrics (as we show in the Results), raising concerns about model generalizability to novel compounds or species. Moreover, while 3D structural fingerprints and advanced machine learning architectures have shown promise, their performance often diminishes for specific classes of compounds. Incorporating taxonomic and physiological information has been shown to improve prediction accuracy, but the extent of this benefit depends heavily on how such information is represented and integrated.

To address these gaps and support rigorous benchmarking of structure-based and species-aware toxicity models, we curated a comprehensive aquatic toxicity benchmark dataset. We focus on aquatic toxicity because it is the most widely available toxicity data. The benchmark aggregates experimentally measured toxicity values from seven major toxicological databases that include data for environmental toxicity models: Aquatic ECETOC [12], Aquatic Japan MOE [22], Aquatic OASIS [27], ECHA REACH [1], ECOTOX [28], envirotox [9], Open food [2]. Furthermore, each species was annotated with taxonomic information following both ITIS and NCBI standards, and linked to physiological descriptors from the Add-My-Pet (AmP) database [21].

Among the most commonly used quantitative measures of aquatic toxicity are the median lethal concentration (*LC*_50_) and the median effective concentration (*EC*_50_). *LC*_50_ represents the concentration of a chemical predicted to cause death in 50% of a tested aquatic population within a specified period, while *EC*_50_ describes a predicted concentration that induces a sublethal effect, such as immobilization or growth inhibition, in 50% of the population. For aquatic invertebrates, acute toxicity is often assessed using immobilization rather than direct mortality, and therefore the resulting *EC*_50_ based on immobilization is commonly used as a surrogate for *LC*_50_ in these studies. These metrics provide the foundation for regulatory toxicity assessments, such as the U.S. EPA’s classification system, which categorizes chemicals into five toxicity levels: “very highly toxic” (*LC*_50_ *<* 0.1 ppm), “highly toxic”, “moderately toxic”, “slightly toxic” and “practically nontoxic” (*LC*_50_ *>* 100 ppm) based on *LC*_50_ thresholds. Predictive models for toxicity prediction typically focus on regression tasks (e.g., predicting *LC*_50_ as in [30]). Here we showed that classification tasks (e.g., predicting toxicity levels) can be useful on its own as well as beneficial for improving regression performance.

Proper assessment is critical when developing predictive models. A frequently overlooked issue is information leakage, in which future or target information inadvertently appears in the training signal [16]. This can arise through different mechanisms including improper data splitting. When leakage occurs, the reported performance is artificially inflated and does not reflect the model’s true ability to generalize to unseen compounds or species. A study that systematically investigated deep learning models for protein-protein interaction (PPI) prediction showed that overlaps between training and test sets resulting from random splitting lead to strongly overestimated performances (in which models learn solely from sequence similarities and node degrees) [5]. Here we investigated the same problem among predictive models for toxicity prediction using different data splitting strategies. Our results showed that random splitting would result in information leakage and therefore inflated performance evaluation.

Finally, we present a new 3D-structure-based toxicity predictor, building on recent advances in 3D representation learning for chemical predictions [14]. Our method is based on 3DMolMS, a deep learning model for MS/MS spectra prediction of compounds from their 3D conformations [13]. It was shown that the molecular representation learned by 3DMolMS can be adapted to enhance the prediction of chemical properties such as the elution time in the liquid chromatography and the collisional cross section measured by ion mobility spectrometry, which are often used to improve compound identification. Here, we demonstrate that this same molecular representation can be utilized, together with species information, to achieve effective cross-compound and cross-species toxicity prediction.

We emphasize that our models, including the new 3D-structure-based model, are designed for cross-species toxicity prediction. This focus is distinct from that of predictors for specific toxic effects, such as those developed for the Tox21 Data Challenge [15]. The Tox21 challenge provided data on 12,000 compounds across 12 toxic effects (like stress response and nuclear receptor effects) from high-throughput assays, where the neural network-based DeepTox was the top-performing model [18].

By integrating rich chemical, taxonomic, and species-level data, our benchmark supports the development and fair evaluation of predictive models that account for both chemical structure and species-specific traits. Moreover, our novel 3D structure–based deep-learning model, 3DMol-Tox, achieves regression accuracy comparable to the best 2D-structure based model while exhibiting consistently higher classification accuracy. Together, this benchmark and our 3D-structure based predictive model provide important tools for advancing computational approaches capable of accurate, generalizable, and animal-free toxicity prediction across diverse aquatic species relevant for environmental toxicology.

## Materials and Methods

### Toxicity Data Sources

We collected LC_50_ and mortality-based EC_50_ records from seven publicly available toxicological databases. These datasets were selected for their regulatory relevance, chemical and species coverage, and data quality. A total of 237,648 raw entries were aggregated before filtering. The final benchmark includes data from the following sources:

- Aquatic ECETOC (ECETOX) [12]: The ECETOC Aquatic Toxicity database is a curated compilation of high-quality, peer-reviewed aquatic toxicity studies published between 1970 and 2000. It comprises more than 5,460 entries covering approximately 600 chemicals. From this dataset, 4,290 mortality-based EC_50_ records meeting our filtering criteria were extracted for inclusion in the benchmark.
- Aquatic Japan MOE [22]: The ecotoxicity database includes 4,596 test results across 668 industrial chemicals. Tests are conducted under harmonized protocols with a focus on LC_50_ and EC_50_ endpoints in aquatic animals. 1,281 records in this dataset match our selection criteria.
- Aquatic OASIS [27]: The Aquatic OASIS database contains curated ecotoxicological test results used for regulatory modeling and QSAR development. All entries are quality-checked and linked to mechanistically relevant chemical categories, making the dataset highly suitable for predictive toxicology. We extracted 2,136 *LC*_50_ and *EC*_50_ records for aquatic species from this collection.
- ECHA REACH [1]: The REACH database spans a broad range of aquatic taxa, test systems, and exposure conditions, although some variability in reporting formats is present due to its self-reported nature. 38,732 LC_50_ and EC_50_ mortality-based records involving aquatic animals were selected from this collection.
- ECOTOX (US EPA) [28]: One of the most comprehensive ecotoxicology databases globally, maintained by the U.S. EPA. The ECOTOX Knowledgebase provides over 1 million records from peer-reviewed literature and government studies, covering thousands of chemicals and species. As of the 2023 release, the database includes 1,170,687 effect records across 12,031 chemicals and 13,376 species. For our benchmark, 112,273 filtered LC_50_/EC_50_ samples were selected.
- OpenFoodTox [2]: OpenFoodTox contains chemical hazard data for 5,712 substances relevant to the food and feed chain, including pesticides, contaminants, and food additives. From this resource, 886 aquatic LC_50_/EC_50_ records were extracted and included in our benchmark following filtering and standardization.
- EnviroTox Platform [9]: A curated and normalized aquatic toxicity database integrating multiple public sources including ECOTOX, ECHA, and legacy repositories. The latest update (September 2021) includes 80,912 effect records across 4,267 unique chemicals and 1,641 species. Its standardized format and taxonomic annotation make it particularly suited for machine learning applications. All 80,912 samples meeting our criteria were included.

### Data Preprocessing, Cleaning and Integration

To ensure consistency across sources and enhance model compatibility, we implemented a unified data preprocessing pipeline. We retained only experimental records with *LC*_50_ or *EC*_50_ endpoints explicitly associated with mortality outcomes and aquatic animal test species. Exposure durations were restricted to 24, 48, 72, or 96 hours, in accordance with standardized test protocols for fish, algae, and crustaceans as established by OECD guidelines [26, 25]. Entries outside this duration range were excluded to minimize biological heterogeneity and align with common regulatory assessment frameworks.

All effect values were converted to a common unit (mg/L) using a controlled keyword-matching strategy that resolved variants such as “ug total silver/L” and “mg a.i./L.” Non-convertible or ambiguously annotated records were discarded. For samples lacking water solubility data, we removed values outside the range of 10^−5^ to 10^5^ mg/L. For entries with solubility data, we also removed effect values that exceeded five times the reported water solubility of the compound. These filtering steps help mitigate potential unit conversion errors, formulation inconsistencies, and misreported test conditions, as similarly noted in prior work [29].

Where available, pH metadata was used to exclude experiments conducted under unrealistic conditions (pH *<* 4 or *>* 10). To address taxonomic inconsistencies, we extracted species-level annotations from each dataset and constructed a canonical taxonomy table by merging all entries and prioritizing authoritative labels from EnviroTox.

To identify erroneous or noisy data points, we applied multiple quality filters. First, we excluded samples exhibiting high variability relative to their mean effect value by applying a standard deviation-to-mean ratio threshold: any group with a standard deviation exceeding 1.5 times the mean was discarded. Next, outliers were removed using a 1.5 × IQR rule within each chemical–species–duration group, provided that at least five replicates were available. To further safeguard against over-filtering, we implemented a cross-species support criterion: extreme outliers were retained if corroborating measurements existed for the same chemical tested on a different species within the same taxonomic order and within a realistic effect value range (10^−5^–10^5^ mg/L). This rule ensures that valuable but rare measurements are preserved when supported by orthologous evidence.

After preprocessing, all cleaned datasets were merged into a unified toxicity matrix. Duplicate entries (defined by CAS number, species name, and exposure duration) were consolidated by averaging effect values and retaining the first non-null metadata fields.

### Handling Missing Taxonomy Using a Hierarchical Inference Procedure

The benchmark contains toxicity data from 2,285 species. Major taxonomic groups including Actinopteri (ray-finned fishes), Amphibia, Branchiopoda, Hexanauplia, Insecta, Malacostraca, and Ostracoda.

To address inconsistencies and missing values in the original species-level taxonomic annotations, we adopted a structured curation strategy centered on preserving the taxonomy as provided by the source databases. The primary taxonomy fields were extracted directly from each source, which largely followed the Integrated Taxonomic Information System (ITIS) conventions. Within this dataset, 1,712 records (corresponding to 392 species) had fully missing taxonomy across all ranks, typically due to unclassified organisms or vernacular names lacking a clear taxonomic map (e.g., “common carp”).

For partially annotated entries, we applied a hierarchical inference procedure to recover as much structured information as possible. Specifically, missing ranks were filled by propagating known lineage information from higher to lower taxonomic levels (e.g., using a known phylum to infer class) and, where appropriate, from lower to higher ranks. If intermediate ranks remained undefined, we assigned fallback labels based on available superclass or broader grouping information, such as using “Unknown class fish” for records categorized only at the superclass Actinopteri. This procedure ensured that major phylogenetic groups (fish, amphibians, and invertebrates) could be consistently distinguished.

This approach balances the retention of a maximum number of records with the need for consistent categorical encoding of taxonomic features, all while preserving the provenance and conventions of the original data sources.

### Machine Learning Models and Train–Test Data Splitting Strategies

Here we evaluated predictive models for aquatic toxicity modeling focus on both regression tasks and classification tasks. For *regression* tasks, the goal is to predict log-transformed LC_50_ values for input chemical compounds. For *classification* task, the goal is to assign compounds into five toxicity levels based on LC_50_ thresholds. Because effect values span several orders of magnitude, the classification formulation provides a complementary view that reflects regulatory decision boundaries and practical toxicity levels.

#### Machine learning models and parameter optimization

We benchmarked three ensemble-based machine learning algorithms that have consistently demonstrated strong and stable performance in quantitative structure–activity relationship (QSAR) modeling: Random Forest (RF), Gradient Boosting (GB), and Gaussian Process Boosting (GPBoost) [31]. All models were trained using a *nested cross-validation* strategy consisting of a randomized hyperparameter search (`n_iter`=3–20, depending on dataset size) with an inner 2- or 3-fold cross-validation, followed by evaluation on an independent test set defined by each data-splitting strategy. This design prevents overfitting to the test set, ensures reproducibility, and provides a robust estimate of model variability across random initializations.

- Random Forest (RF). RF constructs multiple decision trees using bootstrap aggregation and averages their outputs to reduce variance. The following hyperparameters were tuned: maximum tree depth (12–30), number of trees (`n_estimators`=200–800), maximum features per split (`“sqrt”` or `“log2”`), and minimum samples per leaf (1–4).
- Gradient Boosting (GB). GB sequentially fits shallow decision trees to residuals from previous iterations, enabling flexible nonlinear modeling of structure–activity relationships. Tuned parameters included the number of boosting stages (`n_estimators`=100–400), learning rate (0.03–0.2), maximum tree depth (2–4), and subsampling rate (0.8–1.0).
- Gaussian Process Boosting (GPBoost). GPBoost integrates gradient tree boosting with an optional Gaussian process layer to capture structured residual dependencies among observations. In this study, it was used as a LightGBM-style baseline with tuned hyperparameters including learning rate (0.03–0.1), number of leaves (31–63), and maximum depth (3–6).

For all algorithms, categorical taxonomy variables were one-hot encoded, numerical metadata (e.g., exposure duration) were imputed with constant values, and molecular fingerprints (Morgan, MACCS, Mordred, and RDKit; see next subsection) were used as standardized numerical representations of chemical structure. The resulting feature matrices served as input for both regression and classification tasks.

#### Train–test data splitting strategies

To rigorously evaluate model generalization under different structural assumptions, we applied three distinct train–test splitting strategies, summarized below. These strategies progressively reduce structural overlap between training and test compounds and are used consistently throughout all benchmark experiments.

- Random split (“random”): compounds were assigned randomly to training and test sets without considering structural similarity. This setting provides an optimistic baseline but may inflate performance due to information leakage.
- CAS-based group split (“group”): compounds were grouped by CAS Registry Number or equivalent identifiers before splitting, ensuring that related compounds (e.g., analogs or industrial mixtures) appear in only one partition. This simulates realistic regulatory scenarios where structurally related compounds are evaluated together.
- Bemis–Murcko scaffold split (“scaffold”): compounds were partitioned according to their core Bemis– Murcko scaffolds [3], extracted using the RDKit Murcko algorithm. All molecules sharing a scaffold were assigned exclusively to either training or test sets, enforcing strong structural independence and testing extrapolative performance.

For both the group and scaffold splits, a greedy assignment strategy was used to preserve group integrity while maintaining an approximate 80/20 train–test ratio. Table 1 summarizes the dataset sizes for each splitting scheme. These structure-aware splits form the foundation for all subsequent analyses of model generalization and bias mitigation.

**Table 1.** Sizes of the train/test splits under different strategies. All splits cover the same total of 50,343 compounds; small deviations from exactly 80/20 arise from group/scaffold constraints.

| Split | Train (n) | Test (n) | Total | Train % | Test % |
| --- | --- | --- | --- | --- | --- |
| Random | 40,269 | 10,074 | 50,343 | 79.99% | 20.01% |
| Group | 40,292 | 10,051 | 50,343 | 80.03% | 19.97% |
| Scaffold | 40,258 | 10,085 | 50,343 | 79.97% | 20.03% |

### Molecular Fingerprinting Approaches based on 2D Structures

Many different fingerprinting approaches have been developed to represent chemical compounds [10]. To enable benchmarking across different molecular representations, we selected four representative types of chemical descriptors for each unique compound in the dataset: Morgan fingerprints [23], MACCS keys [11], RDKit path-based fingerprints [4], and Mordred descriptors [24]. All structures were standardized using their canonical SMILES (Simplified Molecular Input Line Entry System).

Morgan fingerprints (ECFP4) were generated using RDKit with a radius of 2 and a bit length of 2048. MACCS keys, a rule-based substructure fingerprint, were computed as fixed-length 167-bit vectors. RDKit path-based fingerprints were also generated using a 1024-bit representation. These fingerprints capture complementary structural information ranging from hashed substructures to predefined chemical motifs.

Additionally, we computed 1,613 two-dimensional molecular descriptors using the Mordred descriptor calculator, covering a broad spectrum of chemical properties including topological indices, physicochemical constants, constitutional descriptors, and charge-related features. To ensure consistency in descriptor dimensionality across compounds, descriptor values that could not be computed—typically due to invalid SMILES strings, missing structural elements, or unsupported atom types—were initially encoded as NaN. We then removed all descriptor columns with more than 10% missing values, resulting in a final set of 1,110 descriptors. The remaining missing values were imputed with zero, reflecting the absence of the corresponding structural or physicochemical feature in the compound. This treatment preserves interpretability, as a zero value can be meaningfully interpreted as the non-existence or inapplicability of a particular descriptor. These comprehensive descriptors enable systematic evaluation and comparison of fingerprint types in downstream modeling tasks.

### 3DMol-Tox: A Predictive Model using 3D-structure based Fingerprints

3D structure-based fingerprints have been leveraged to provide better solutions to problems including chemical property prediction [17] and compound identification from their MS/MS spectra as in 3DMolMS [13]. Here we test if 3D structure-based fingerprints can be leveraged for toxicity prediction. Similar to 3DmolMS [13], 3DMol-Tox utilizes RDkit to generate 3D structures from SMILES. As in 3DmolMS [13], 3DMol-Tox uses a graph neural network (see Figure 1) for toxicity prediction given 3D coordinates of chemical compounds.

**Figure 1.**
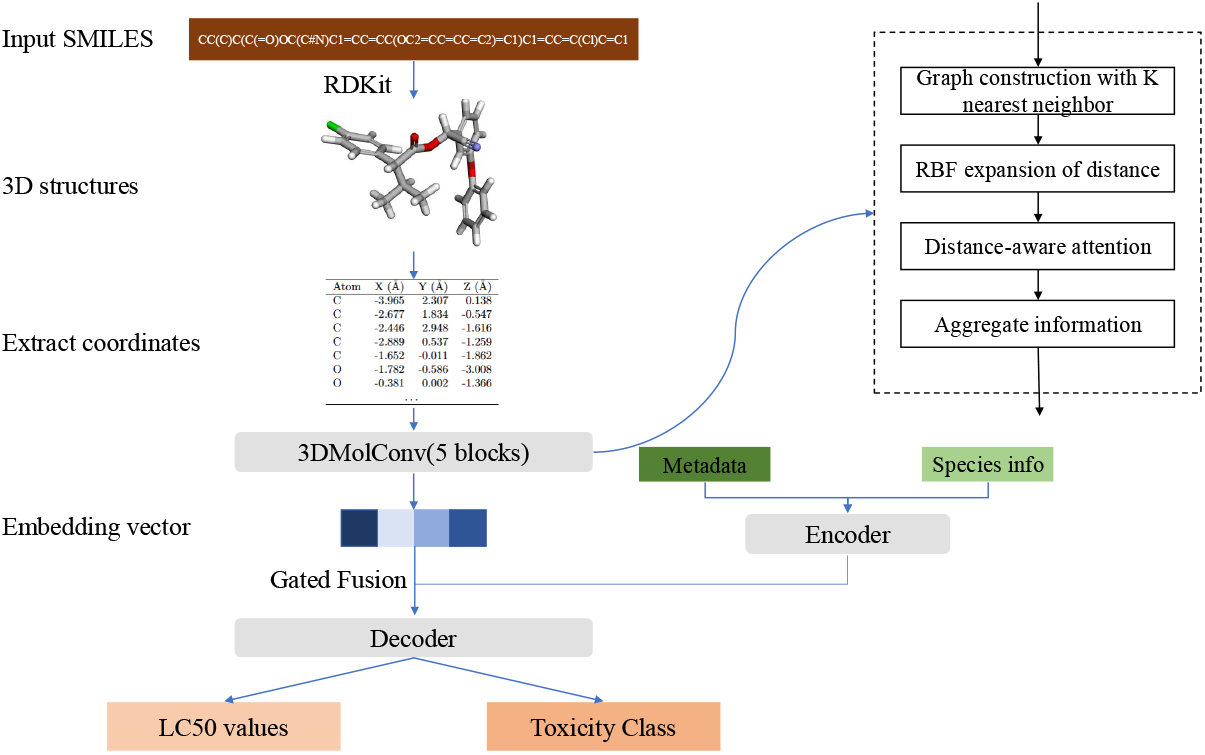
The 3DMol-Tox model architecture. 3DMol-Tox is based on the 3DMolMS, a method for prediction of mass spectra from 3D structures. 3DMol-Tox incorporates the embedding of chemical compounds and the species representation (deb or taxonomy) for prediction of toxicity.

#### Input representation and preprocessing

For each molecule, we generate up to three low-energy conformers. Each atom is described by a feature vector comprising: (i) 3D coordinates (*x, y, z*) and (ii) categorical descriptors (e.g., atom-type one-hots from a predefined vocabulary). We cap the atom count at 300 and zero-pad shorter molecules. Training uses only one conformer for validation/testing and multiple conformers for augmentation during training.

#### Graph construction

Given coordinates *X* ∈ ℝ^*N* ×3^, we connect each atom *i* to its *k* nearest neighbors by Euclidean distance, yielding a directed local *k*-NN graph. Distances *d*_*ij*_ = ∥ **x**_*i*_ − **x**_*j*_ ∥ _2_ are encoded via a radial basis expansion

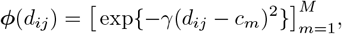

with *M* kernels, centers {*c*_*m*_} uniformly spanning a cutoff window, and bandwidth *γ*. This preserves translation- and rotation-invariance at the message level, since messages condition on inter-atomic distances rather than raw coordinates.

#### MolConv with distance-conditioned attention

We replace the original distance/Gram gating from 3DMolMS [13] with a lightweight RBF-conditioned attention that scores each neighbor using both chemistry and geometry. For a central atom *i* and a neighbor *j*, let **h**_*i*_, **h**_*j*_ ∈ ℝ^*C*^ be their current features. We form the concatenated token

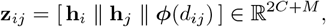

and compute attention logits with a 1×1 Conv–MLP:

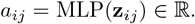

Attention weights are *α*_*ij*_ = softmax_*j*∈𝒩*i*)_(*a*_*ij*_), and the layer aggregates neighbor features by

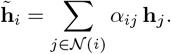

An update MLP (pointwise 1 × 1 convolution, BatchNorm, and LeakyReLU) maps 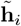 to the next feature space. We then average over the *k* neighbors. Compared to the original gating using distance and double-Gram matrices, the RBF-attention (i) learns non-linear distance sensitivity, (ii) fuses geometry with atom identity directly in the attention MLP, and (iii) removes the *k*×*k* Gram construction cost.

#### Encoder–decoder architecture

3DMol-Tox employs a lighter but more stable encoder–decoder design. The encoder consists of five MolConv blocks with hidden widths {64, 64, 128, 256, 512}, progressively expanding the local receptive field to aggregate geometric and chemical information. A permutation-invariant pooling layer produces a molecular embedding of dimension 1024. The decoder is a shallow three-layer MLP {1024, 256, 64} followed by two task heads for regression and ordinal classification. A dropout rate of 0.3 is applied throughout the decoder to regularize training and improve generalization. Each molecule is truncated or zero-padded to a maximum of 300 atoms, and species-level or environmental embeddings (dimension 256) are concatenated to the pooled molecular vector before decoding.

#### Multi-task objective

3DMol-Tox jointly optimizes three complementary objectives: (i) regression of log_10_(LC_50_) via mean-squared error (ℒ_reg_); (ii) multi-class toxicity classification using cross-entropy (ℒ_ce_); and (iii) cumulative ordinal regression with logistic (CORAL [6]) thresholds (ℒ_coral_), equipped with adaptive per-threshold weights to mitigate class imbalance. The total loss is a weighted sum:

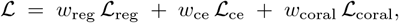

This joint formulation balances continuous and discrete toxicity learning, stabilizes convergence, and encourages smooth ordinal transitions between regulatory toxicity categories.

#### Optimization and training schedule

Models were trained for 100 epochs using AdamW (learning rate 10^−3^, batch size 32) with early stopping based on validation loss. Cosine or plateau-based learning-rate decay was applied for fine convergence. The MolConv neighborhood size was fixed to *k*=5, with *M* =16 radial basis kernels, *γ*=10. All experiments used consistent random seeds and the same preprocessing pipeline to ensure comparability across runs.

### Metrics for Performance Evaluation

We used a variety of metrics to evaluate the performance of predictive models.

- **Root Mean Squared Error (RMSE)** quantifies the average magnitude of prediction errors in the log-transformed LC50 values. Although commonly used in regression, its biological interpretation can be limited, as large errors in extremely high or low toxicity values (e.g., between 0.00001 and 0.1 mg/L) may have minimal impact on toxicity categorization but disproportionately affect RMSE.
- **R Squared (coefficient of determination)** measures the proportion of variance in the true LC50 values explained by the model. While useful for gauging overall model fit, high *R*^2^ does not guarantee correct toxicity classification, especially near decision thresholds.
- **Accuracy** measures the proportion of correctly classified samples among all predictions. While intuitive, it treats all misclassifications equally—predicting a *very highly toxic* compound as *practically nontoxic* is penalized the same as misclassifying it as *highly toxic*, despite vastly different biological implications.
- **Weighted F1-score** balances precision and recall across classes, accounting for class imbalance. It is particularly important in toxicity classification, where underrepresented classes (e.g., *highly toxic*) are clinically significant and may be overlooked by accuracy alone.
- **Within-One-Bin Accuracy (W1B)** For ordinal toxicity labels, small near-misses are often acceptable. W1B counts a prediction as correct if it is in the true bin or an adjacent bin:

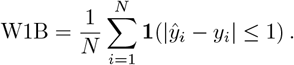

For example, if the true class is *moderately toxic* (bin 3), predictions as bins 2, 3, or 4 are all considered accurate. This reflects regulatory tolerance and assay variability.

Together, these complementary metrics allow us to assess not only numerical accuracy but also practical and biological relevance, which is critical for evaluating predictive models intended to guide chemical safety decisions.

## Availability

The proposed Python library tox-learn, including the benchmarks, machine learning methods, the different train-test splittings, and performance evaluation are publicly available at https://github.com/mgtools/tox-learn. The repository includes the source code, instructions for installation, and guidelines for using the library. The new model 3DMol-Tox is also available in the same repository.

## Results

### Benchmark Statistics and Chemical Composition Distribution

Application of our data integration process yielded a final benchmark comprising 50,603 records (of 5,889 compounds involving 2,285 species) after removing 7,412 outliers and consolidating overlapping records. For comparison, the dataset used in benchmarking Bio-QSAR 2.0 contains approximately 20,000 samples [31].

Figure 2A shows the distribution of the effect values of the compounds included in our benchmark. These compounds were grouped into five toxicity levels. As Figure 2B shows, the benchmark exhibits a moderately balanced composition, with the *slightly toxic* (24.2%) and *moderately toxic* (24.0%) classes being most frequent, and the *highly toxic* (17.4%) and *practically nontoxic* (15.8%) classes somewhat underrepresented.

**Figure 2.**
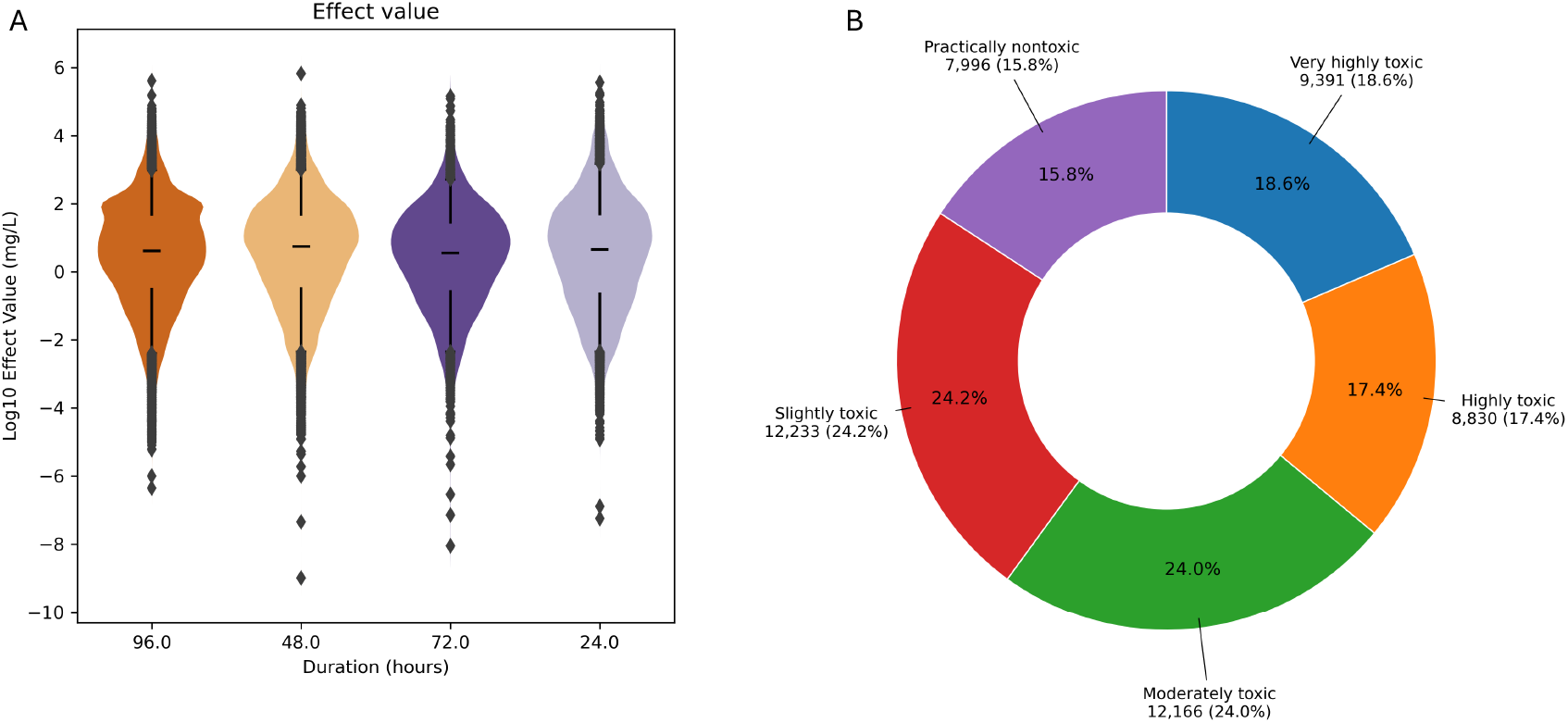
Distribution of toxicity effect values and categories. (A) shows the distribution of the effect values of the chemical compounds included in the toxicity benchmark in violin plots. (B) shows the overall distribution of LC_50_-based toxicity categories across all compounds and species.

To assess the chemical diversity of the benchmark dataset, we performed a comprehensive analysis of elemental composition and functional group occurrence across all unique molecular structures. Each compound was assigned to one of five broad chemical categories—organic, organometallic, inorganic metal, inorganic nonmetal, or unknown—based on the presence or absence of carbon and metallic elements. Functional groups were identified using SMARTS-based substructure matching implemented in RDKit, covering major classes such as halogens, amines, esters, amides, and heteroaromatic rings.

Figure 3 summarizes the overall distribution of chemical categories and functional groups. The dataset is overwhelmingly composed of organic compounds (91.9%), with smaller fractions of inorganic nonmetals (4.6%), organometallic species (1.8%), and inorganic metals (1.6%). Among functional groups, halogenated (19.5%) and heteroaromatic (10.6%) structures are most prevalent, followed by amines, esters, and amides (each 6–10%). These results indicate that the benchmark predominantly consists of small organic molecules bearing diverse heteroatomic and polar functionalities, characteristic of industrial and environmental contaminants commonly represented in regulatory ecotoxicological databases.

**Figure 3.**
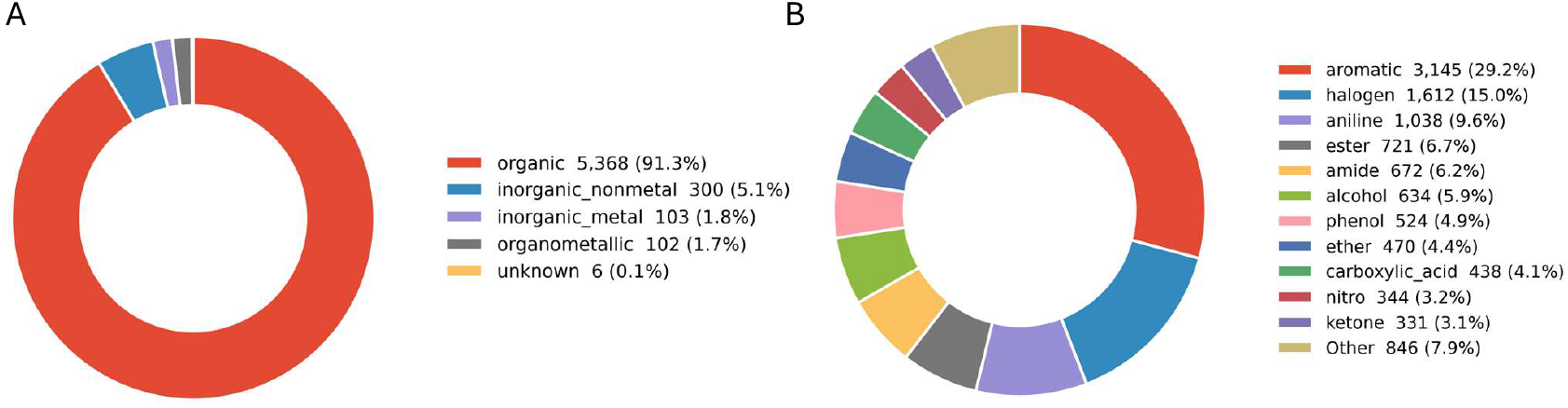
Chemical composition of the unified benchmark dataset. (A) Broad chemical categories. (B) Major functional groups. Pie charts were generated using RDKit-based SMARTS substructure detection and Matplotlib visualization.

### Evaluating the Impact of Model, Fingerprint, and Splitting Strategy

To evaluate how model architecture, molecular representation, and data-splitting strategy jointly influence predictive performance, we benchmarked all combinations of three ensemble-based methods (RF, GB, and GPBoost) and four fingerprint types (Morgan, MACCS, Mordred, and RDKit) under the three splitting protocols (random, group, and scaffold). Figure 4 illustrates the distribution of the similarity between the test and train subsets generated by each of these splitting protocols.

**Figure 4.**
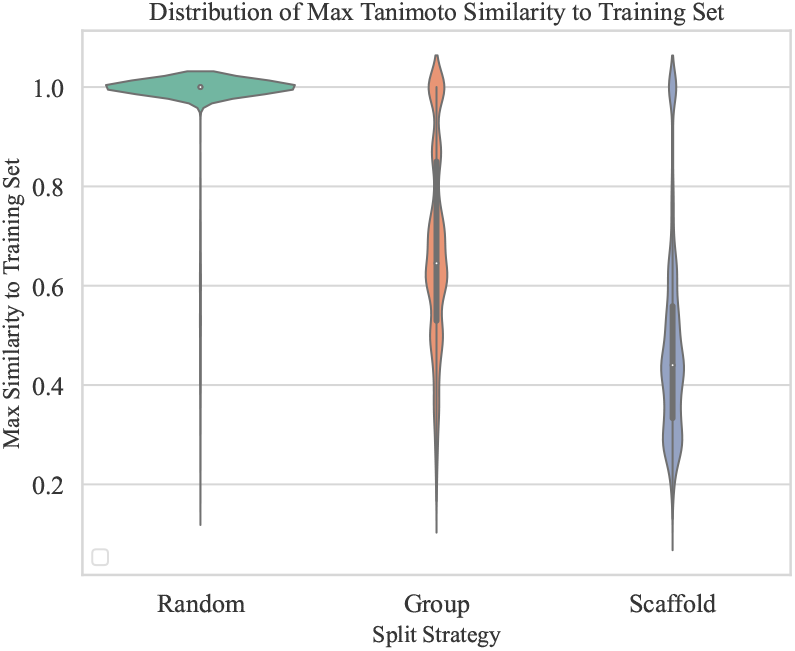
Violin plots showing the distribution of maximum Tanimoto similarity (based on Morgan fingerprints, radius=2) between test and train compounds, with random split showing the highest structural similarity and scaffold split the lowest.

A model was trained for each configuration using the nested cross-validation and hyperparameter optimization procedures described in the Methods section. Table 2 summarizes regression and classification results across all model–fingerprint–split combinations. Figure 5A-C show how different fingerprinting tools and models perform under three data splitting methods. It reveals how certain model-fingerprint combination significantly dominate within specific split while some remain statistically indistinguishable. Several consistent trends emerged:

**Table 2.**
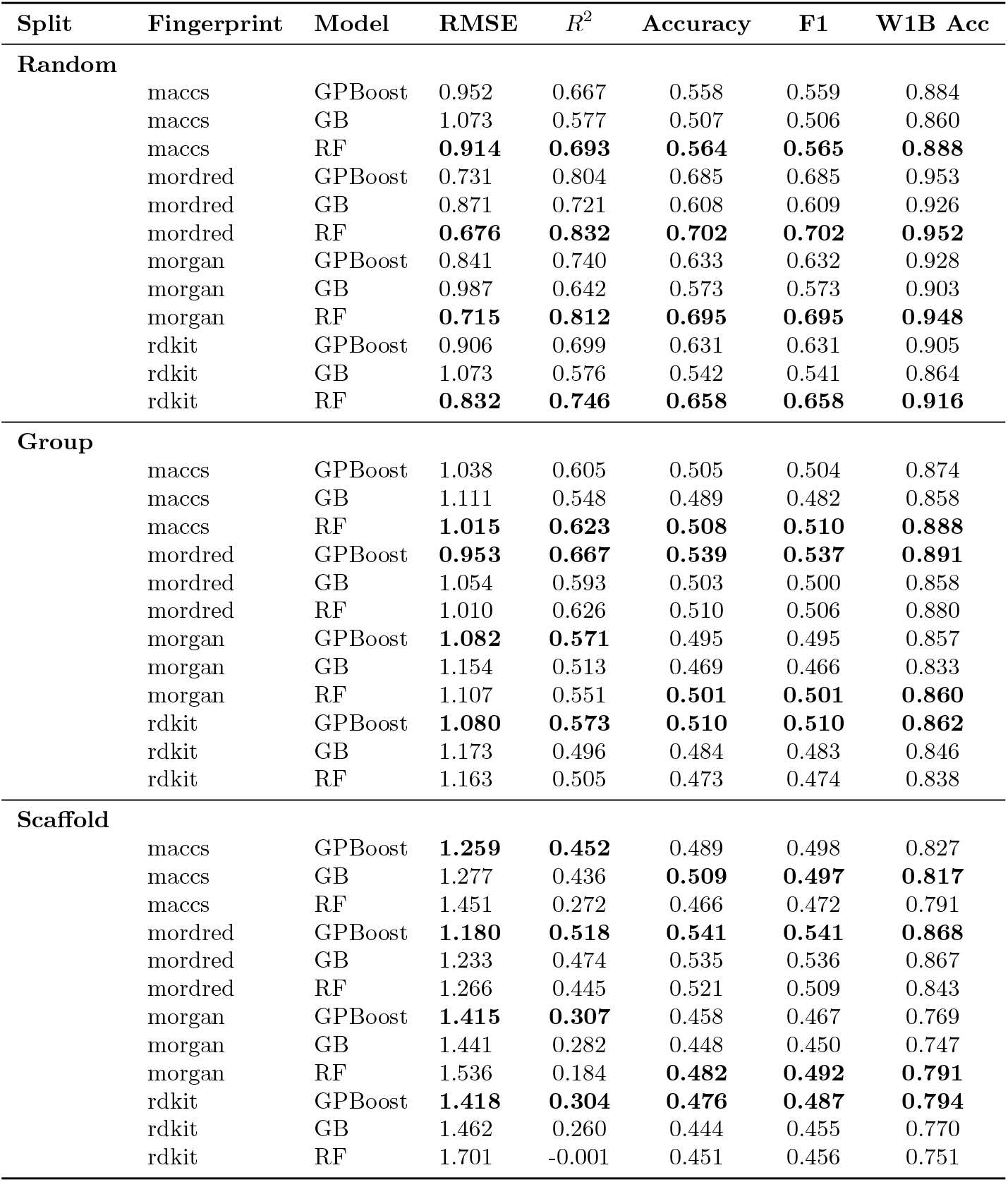
Comparison of regression and classification performance across splits, fingerprints, and models (with lower RMSE, higher *R*^2^, accuracy, F1, and W1B accuracy indicating better performance; boldface highlights best results for each fingerprint–split configuration).

**Figure 5.**
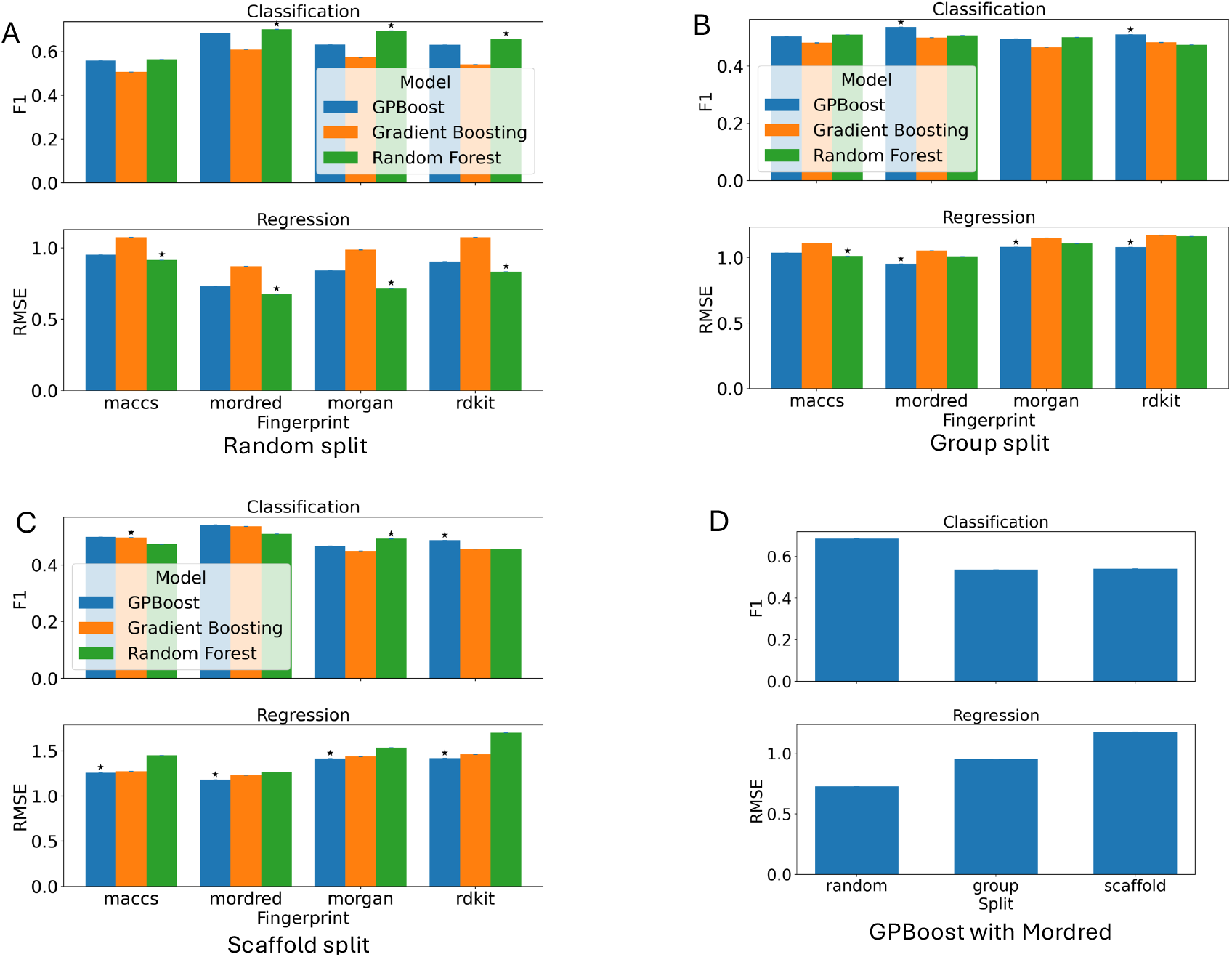
Comparison of the performance by the different predictive models when different fingerprints were used. Panels A-C show the comparison of the models when random, group-based, and scaffold-based train-test splitting were applied, respectively. Stars on top of the bars indicate the models that significantly outperformed the other two according to pairwise bootstrap tests (*p* < 0.05). Panel D shows the comparison of the performance of GPBoost using Mordred fingerprints across different train-test splitting schemes, showing the drastic drop of the performance (reflected by lower F1 score and higher RMSE) when information leakage was reduced by the group-based and scaffold-based train-test splitting.

- Under the random split, models benefited from structural overlap between training and test sets, yielding the highest apparent performance. RF combined with Mordred fingerprints achieved the lowest regression error (RMSE = 0.676) and highest classification accuracy (0.702), reflecting its capacity to exploit detailed physicochemical descriptors when information leakage is present.
- The group split, which preserves related compounds while minimizing exact duplication, provided a more realistic measure of generalization. Here, GPBoost + Mordred achieved the best balance between regression and classification (RMSE = 0.953, Accuracy = 0.539), slightly outperforming RF and GB. This configuration most closely approximates real-world ecotoxicological screening, where new species or conditions are encountered for structurally similar compounds.
- The scaffold split imposed the strictest constraint by excluding shared molecular frameworks between training and test compounds. As expected, overall performance decreased across all methods, illustrating the difficulty of extrapolating toxicity to unseen scaffolds. GPBoost + Mordred again yielded the strongest regression result (RMSE = 1.180) and maintained competitive classification accuracy (0.541), demonstrating robust extrapolative capability.
- Across all experiments, Mordred fingerprints consistently outperformed Morgan, MACCS, and RDKit fingerprints. Their advantage likely arises from richer physicochemical and topological descriptors, which enable the models to capture subtle structure–toxicity relationships beyond binary substructural matches.

Figure 5D shows the performance of GPBoost using the Modred fingerprint under different data-splitting strategies. Performance dropped substantially when information leakage was minimized by using group-based or scaffold-based train–test splits. As shown in Figure 4, compounds in the test subset shared high structural similarity with those in the training subset under random splitting, leading to inflated performance estimates. Therefore, results based on random train–test splitting should be taken with reservation.

Collectively, these findings reveal three key insights: (i) random splits substantially overestimate generalization due to scaffold overlap; (ii) group splits provide the most realistic and reproducible benchmark for practical toxicological applications; and (iii) scaffold-based evaluations remain indispensable for assessing true extrapolation to novel chemical space. We therefore identify the group split with Mordred fingerprints as the most balanced and interpretable configuration for cross-chemical and cross-species toxicity prediction.

### Incorporation of Species Information Improves Performance

To evaluate the impact of species-level information on QSAR model performance, we benchmarked four different species representations: (1) the baseline one-hot encoding of species identity (`ablation`), (2) taxonomy from the original dataset (`original`), (3) standardized hierarchical taxonomy from NCBI (ncbi), and physiological traits derived from Dynamic Energy Budget (DEB) theory (`deb`) [20]. The last representation was used in combination with chemical fingerprints by Bio-QSAR for cross-chemical and cross-species toxicity prediction [30].

For this analysis, we trained a GPBoost model using Mordred fingerprints on the data subset where all three types of species information overlapped. This resulted in 24,672 training samples and 5,367 test samples. We evaluated model performance on two tasks, regression (log-transformed LC_50_) and multiclass classification (EPA toxicity categories), using the three different data splitting strategies, respectively. Table 3 summarizes the results.

**Table 3.** Evaluation of the effects of different species representations on predictive models under various splitting strategies (with lower RMSE, higher *R*^2^, accuracy, F1, and W1B accuracy indicating better performance; boldface highlighs best results for each split).

| Split | Dataset | RMSE | $R^2$ | Accuracy | F1 | W1B Acc |
| --- | --- | --- | --- | --- | --- | --- |
| Random | ablation | 0.660 | 0.826 | **0.723** | **0.723** | **0.955** |
|  | original | 0.654 | 0.829 | 0.706 | 0.706 | 0.948 |
|  | ncbi | 0.649 | **0.832** | 0.710 | 0.710 | 0.949 |
|  | deb | **0.648** | **0.832** | 0.708 | 0.708 | 0.947 |
| Group | ablation | 0.977 | 0.654 | 0.594 | 0.596 | 0.885 |
|  | original | 0.822 | **0.755** | 0.629 | 0.630 | 0.897 |
|  | ncbi | 0.823 | 0.755 | 0.630 | 0.631 | 0.898 |
|  | deb | **0.822** | **0.755** | **0.633** | **0.635** | **0.901** |
| Scaffold | ablation | 1.144 | 0.535 | 0.506 | 0.512 | 0.842 |
|  | original | **1.085** | 0.581 | **0.522** | 0.514 | **0.851** |
|  | ncbi | 1.100 | **0.570** | 0.519 | **0.518** | 0.848 |
|  | deb | 1.133 | 0.543 | 0.521 | 0.512 | 0.849 |

As expected, model performance was highest under random splits, which permit structural redundancy between training and test sets. Regression RMSEs reached as low as 0.648 with *R*^2^ values exceeding 0.83, and classification accuracy exceeded 0.70. However, these metrics are likely overestimates of true generalization ability due to information leakage. Scaffold splits, which enforce structural disjointness, induced substantial performance degradation. Group splits, which stratify by CAS identifiers and control for both species and chemical duplication, yielded intermediate performance while better reflecting prospective regulatory use cases.

Biologically meaningful species encodings consistently outperformed naïve baselines. Under random splits, `deb` and `ncbi` achieved the strongest regression performance (*R*^2^ ≈ 0.83), underscoring the value of mechanistic and phylogenetic descriptors. Interestingly, the `ablation` baseline achieved the highest classification accuracy (0.723), likely due to memorization of species-specific labels in the absence of generalization pressure. This advantage disappeared under scaffold and group splits, where `ablation` performance deteriorated markedly (e.g., scaffold RMSE = 1.144, *R*^2^ = 0.535; accuracy = 0.506).

In contrast, both `deb` and `ncbi` demonstrated stable performance across all splits. Under scaffold split, ncbi achieved the highest regression *R*^2^ (0.570), and under group split, `deb` outperformed all others in both regression (*R*^2^ = 0.755) and classification (accuracy = 0.634, F1 = 0.635). The original taxonomy representation (`origin`) closely followed these two but lacked their robustness to data perturbation.

These results emphasize the critical role of species-level biological context in modeling interspecies toxicity. The superior performance of deb and ncbi representations stems from their incorporation of biological function (e.g., metabolic rate, reproduction) and evolutionary relationships, respectively—factors known to modulate toxicodynamic responses in ecological species. The poor generalization of `ablation` further highlights the risk of overfitting when using species ID as a proxy without underlying biological descriptors.

Ultimately, these findings suggest that effective multi-species QSAR modeling requires biologically grounded features. Combining mechanistic physiology (deb) and phylogenetic hierarchy (ncbi) provides a path forward for robust toxicity prediction across taxonomic diversity.

### Evaluation of 3DMol-Tox Performance

We evaluated the performance of 3DMol-Tox and compared it to GPBoost. 3DMol-Tox relies on molecular conformers generated by RDKit. However, RDKit often fails to produce valid conformers for compounds containing salts, mixtures, or charged complexes (e.g., Na^+^, Cl^−^). Therefore, all compounds with ions or disconnected fragments were excluded from this analysis. For each remaining molecule, a single lowest-energy conformer was generated, from which 3D atomic coordinates and geometric features were extracted.

A key difference in our approach is that a single 3DMol-Tox model was trained to handle both classification and regression tasks simultaneously. For comparison, separate GPBoost models were trained for classification and regression, respectively. To capture the ordinal nature of the toxicity levels, 3DMol-Tox employs the CORAL (Cumulative Ordinal Regression with Logistic link) approach [6], which imposes a monotonic constraint on class boundaries and improves ordinal consistency.

For statistical robustness, we generated 10 independent dataset replicates using different random seeds for data splitting, ensuring that results reflect performance consistency rather than a single partition. The mean values across these ten runs are reported in Table 4. Across both group and scaffold splits, 3DMol-Tox achieves regression accuracy comparable to GPBoost while exhibiting consistently higher within-one-bin (W1B) accuracy, suggesting that using geometric information (when valid 3D conformers are available) and multi-tasking learning enhances ordinal smoothness across toxicity categories.

**Table 4.** Benchmark comparison between GPBoost and 3DMol-Tox on group and scaffold splits (boldface indicates the best performance for each metric).

| Split | Model | RMSE | $R^2$ | Accuracy | F1 | W1B Accuracy |
| --- | --- | --- | --- | --- | --- | --- |
| Group | GPBoost | 0.978 | 0.650 | **0.534** | **0.533** | 0.881 |
|  | 3DMol-Tox | **0.952** | **0.674** | 0.527 | 0.525 | **0.917** |
| Scaffold | GPBoost | 1.144 | 0.539 | 0.501 | 0.502 | 0.864 |
|  | 3DMol-Tox | **1.080** | **0.571** | **0.522** | **0.517** | **0.891** |

### Analysis of Classification Errors in Chemical Space

To compare the performance of 2D-structure based GPBoost model and 3DMol-Tox across chemical space, and to identify where the each model performed well or poorly, we proposed a compound-level correctness score (*p*_W1B_) and applied a dimensionality reduction to project chemical compounds in lower dimensional space (2D) for visualization.

Specifically, *p*_W1B_ is the compound-level within-one-bin accuracy, which is the fraction of test samples (tested in different species) for that compound whose predicted toxicity category falls within one ordinal bin of the true label. For visualization, compounds in the test samples from the scaffold splitting were represented using Mordred fingerprints, which were standardized and then reduced to two dimensions using t-distributed stochastic neighbor embedding (t-SNE). In addition, KMeans clustering (k=64) was applied to the standardized Mordred fingerprints to group compounds to facilitate the analysis.

Figure 6 shows the test compounds in the 2D chemical space, colored by toxicity prediction accuracy (*p*_W1B_) for GPBoost (left) and 3DMol-Tox (right). Each point represents a test molecule, with warmer tones indicating lower correctness. Overall, both models achieved broadly high accuracy across the chemical space, performing well on some compounds (green points in both plots) and poorly on others (warm-colored points). However, notable differences were found: 3DMol-Tox resulted in more accurate predictions as a whole (W1B of 0.891 versus 0.864; see Table 4) and differential performance patterns were evident for certain groups of compounds, such as those highlighted in Figure 6.

**Figure 6.**
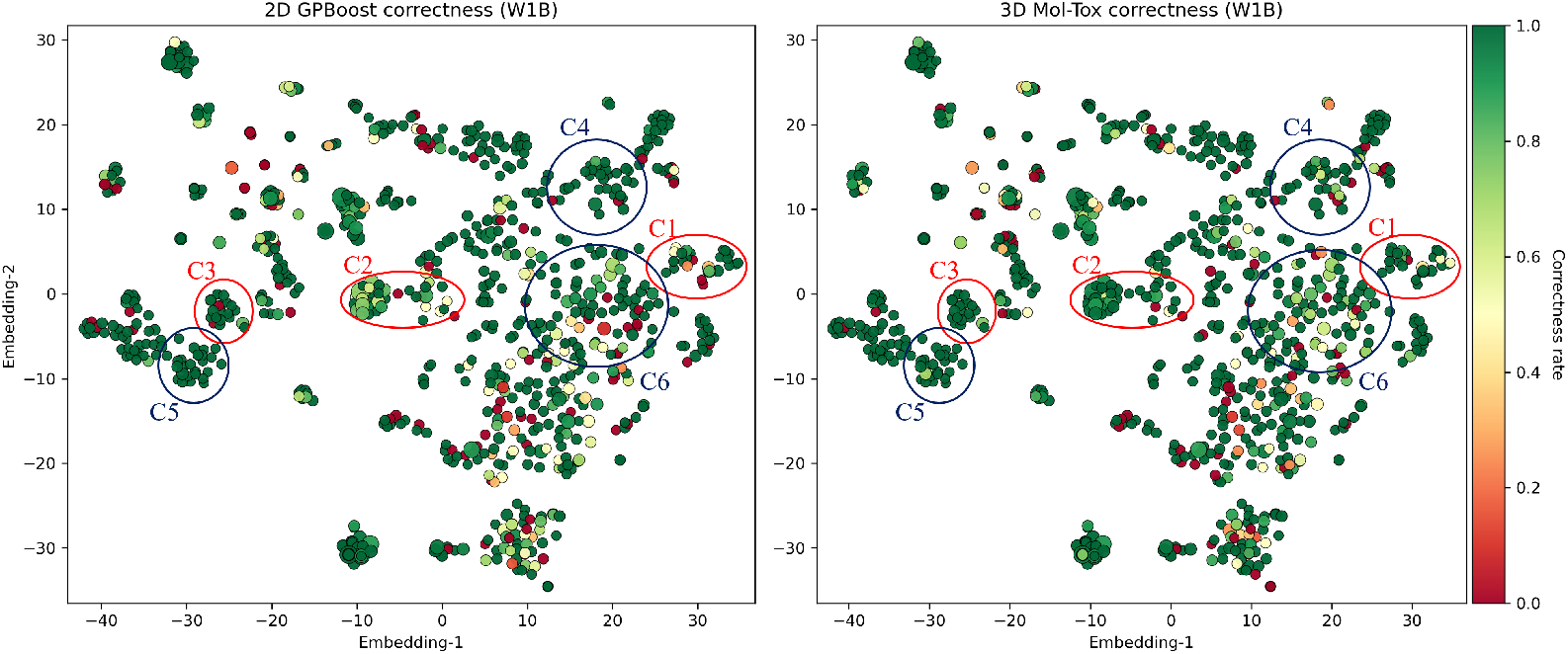
Visualization of compounds in the chemical space to show how well 2D-structure based model (GPBoost) and 3D-structure based model (3DMol-Tox) performed. Each point represents a test compound colored by compound-level prediction accuracy (*p*_W1B_) (cold colors indicate more accurate predictions). Red ellipses highlight a few clusters of compounds (C1, C2 and C3) where 3DMol-Tox outperformed the 2D model, while blue ellipses mark regions (C4, C5 and C6) where the 2D model performed better.

Table 5 summarizes the average within-one-bin correctness and differences for both models for compounds in these selected groups. Clusters C1-C3 (red ellipse) represent the groups of compounds where 3DMol-Tox had substantial improvements over 2D model (Δ*p* up to +0.19), suggesting that geometric information can be leveraged to improve prediction correctness for certain compound groups. In contrast, for clusters C4-C6 (blue ellipse), although 2D model had slightly higher correctness, the margin is small (Δ*p <* 0.07) and often appear in the regions where 2D model already achieved near perfect accuracy. These results suggest that a hybrid model could be devised to take advantage of the complementary strengths of 2D and 3D fingerprints.

**Table 5.** Representative clusters of compounds with performance differences between 2D and 3D models.

| Cluster | $p_{2D}$ | $p_{3D}$ | $\Delta p$ | samples | compounds | dominant FG (frac.) |
| --- | --- | --- | --- | --- | --- | --- |
| C1 | 0.64 | 0.83 | +0.19 | 130 | 25 | Aromatic (1.00) |
| C2 | 0.75 | 0.93 | +0.18 | 806 | 35 | Aromatic (0.97) |
| C3 | 0.88 | 0.92 | +0.05 | 66 | 23 | Ester (0.26) |
| C4 | 0.96 | 0.89 | -0.07 | 131 | 34 | Aromatic (1.00) |
| C5 | 1.00 | 0.94 | -0.06 | 98 | 34 | Ether (0.35) |
| C6 | 0.84 | 0.79 | -0.05 | 347 | 53 | Aromatic (1.00) |

This table lists the average prediction correctness of the compounds in each cluster by the 2D and 3D models ( $p_{2D}$ and $p_{3D}$ ), their difference $\Delta p = p_{3D} - p_{2D}$ , the number of test samples and unique compounds, and the dominant functional group (FG) in the compounds for each cluster.

## Discussion

This work establishes a rigorous foundation for cross-chemical and cross-species toxicity prediction by unifying data quality control, representation learning, and structure-aware evaluation. Through a comprehensive benchmarking framework encompassing over fifty thousand experimentally measured LC_50_ records, we demonstrate that predictive performance in QSAR modeling is shaped as much by data partitioning and representation design as by algorithmic sophistication.

Our results show that random train–test splits—commonly adopted in prior studies—systematically overestimate model performance due to scaffold and biological redundancy. In contrast, group-, scaffold-, and similarity-based splits reveal the true generalization limits of toxicity models, with performance degrading predictably as structural novelty increases. Among these, the group-based split provides a practical balance between realism and robustness, approximating prospective deployment scenarios in regulatory screening where known chemical series are evaluated across new taxa or experimental contexts.

Based on our comprehensive benchmarking and analysis, we propose the following recommendations for QSAR model evaluation, aligning machine learning practices with biologically meaningful scenarios:

- **Deprecate random splits** as the default benchmarking strategy. While random splits often yield high predictive performance due to scaffold leakage, they reflect an unrealistic scenario where both chemical structure and biological context are highly redundant between training and test sets. However, random splits may serve as a useful reference when evaluating model performance on previously studied chemicals tested in new species, where structural novelty is limited but biological extrapolation is desired.
- **Adopt group-based splitting** as the default evaluation protocol for realistic toxicological applications. Group splits partition data by compound identity (e.g., CAS number), species, and experimental conditions, enabling assessment of model generalization to new species or test environments for structurally related compounds. This mirrors regulatory and screening scenarios where known scaffolds are evaluated across diverse biological systems.
- **Use scaffold-based splits** as a secondary diagnostic tool for assessing extrapolation to novel chemical series. By assigning entire scaffolds to either the training or test set, this strategy rigorously tests the model’s ability to predict toxicity for new scaffolds that differ structurally from any seen during training. It is particularly useful for early-stage drug discovery or chemical design tasks where structurally novel molecules are anticipated.
- **Reserve similarity-based splits** for stress-testing model robustness under extreme distribution shifts. These splits create maximal structural disjointness, with no test compound having high similarity to any training molecule. While this provides insight into extrapolation limits, the overly stringent nature of this split may underestimate model utility in real-world settings and is thus not recommended as a routine benchmarking strategy.

We further highlight that predictive accuracy depends critically on both chemical and biological representations. Mordred fingerprints consistently outperform alternative 2D encodings by capturing a richer spectrum of structural and physicochemical features. Likewise, biologically informed species descriptors—derived from Dynamic Energy Budget theory or standardized phylogenetic hierarchies—yield substantially more transferable models than one-hot encodings of species identity. These findings emphasize that mechanistic and evolutionary context are not ancillary but foundational to cross-species generalization.

Our newly introduced 3DMol-Tox model demonstrates that 3D molecular geometry can enhance ordinal smoothness and local consistency, particularly for heteroaromatic and orientation-dependent functional groups. While its classification accuracy remains comparable to descriptor-based baselines, the improved within-one-bin alignment indicates that integrating geometric features with physicochemical fingerprints offers a promising path toward unified 2D–3D modeling frameworks.

Beyond methodological insight, this study provides a quantitative basis for reproducible, structure-aware benchmarking in computational toxicology. By explicitly measuring structural similarity, partitioning data to avoid information leakage, and incorporating biologically grounded species features, we define best practices that align machine learning evaluation with ecological and regulatory realism.

Looking forward, we envision extending this framework toward multitask and foundation models that jointly learn across endpoints, species, and environmental contexts. A promising direction is the development of a hybrid, region-aware modeling framework that dynamically integrates 2D and 3D representations based on 3D structure information gain. This adaptive fusion of complementary representations could be embedded within larger multitask architectures, enabling simultaneous learning of toxicity, mode-of-action, and cross-species transfer. Coupling structure-aware splitting with interpretable, mechanistic embeddings could enable predictive systems that are not only more accurate but also more transparent and biologically credible—an essential step toward replacing traditional animal assays with trustworthy AI-driven toxicity assessment.

## Notes

https://github.com/mgtools/tox-learn

### Competing Interest Statement

The authors have declared no competing interest.

### Funding

- U.S. National Science Foundation: DBI-2011271

